# Assessing the helical stability of polyXYs at the boundaries of Intrinsically Disordered Regions with MD simulations

**DOI:** 10.1101/2024.11.16.623902

**Authors:** Mariane Gonçalves-Kulik, Luis A. Baptista, Friederike Schmid, Miguel A. Andrade-Navarro

## Abstract

Intrinsically disordered regions (IDRs) in proteins lack stable structure. By carrying many hydrophilic and charged residues, it prevents them from forming globular domains and contributes to their flexibility and accessibility. Naturally, regions with reduced amino acid composition (low complexity regions; LCRs) occur within IDRs. Disorder and low complexity in protein sequences are linked to various biological functions including phase separation, regulation, and molecular interactions, and mutations in these regions can contribute to several diseases including cancer. Understanding these biological properties requires examination of the structural properties of IDRs and the LCRs within, but their inherently dynamic nature requires specific approaches combining sequence analyses, structure predictions and molecular dynamics (MD) simulations. Here, we leverage our previous work where we identified that some types of LCRs combining two residues (polyXY) are frequent within IDRs and confer them with propensity to form helical conformation. We identify a significant accumulation of such polyXYs at the ends of IDRs following alpha-helices that start outside the IDR and may extend through the polyXY into the IDR, particularly from the N-terminal end of the IDR. MD simulations support the dynamic nature of such helical conformations. Our results suggest a mechanism by which evolutionary emergence of LCRs at IDR ends could provide proteins with flexible regions for fold-upon-binding.

## Introduction

Intrinsically disordered regions (IDRs) are regions in protein sequences that do not adopt a stable tertiary structure and often not even secondary structures, remaining flexible and widely accessible by the solvent [1], [2]. These regions are characterized by the accumulation of hydrophilic amino acid residues, making them unable to form the hydrophobic core required for the formation of globular domains. Another important attribute of IDRs is the accumulation of highly charged residues, which are believed to influence the compaction and expansion of these regions [3], [4].

The natural reduction of the amino acid diversity within those regions often propitiates the accumulation of residues with similar physico-chemical properties, resulting in low complexity regions (LCRs). Despite not being restricted to IDRs, these regions are common within them. They can span extremely low complexity, accumulating several copies of the same amino acid, known as homorepeats, to more complex patterns composed of different residues with similar characteristics [5], [6], [7].

Both disorder and low complexity in protein sequences were extensively explored in recent years, due to their biological relevance. Protein regions where these properties associate perform several important biological processes in the cell, ranging from driving liquid-liquid phase separation and the formation of biomolecular condensates, to performing transcriptional and translational regulation, molecular recognition and binding, to mention a few [8], [9], [10]. Mutation in these regions might lead to diseases, including a range of neurodegenerative diseases, such as amyotrophic lateral sclerosis (ALS), Alzheimer’s, Parkinson’s and Huntington’s disease, neurodevelopmental disorders and many types of cancer [11], [12].

Due to their plasticity, experimental resolution of the structures of IDRs is challenging, requiring the application of specific techniques to ensure that their multiple conformations are properly captured. As a result, these regions are underrepresented in structure repositories and our knowledge about their structural dynamics and functional behavior is limited [13], [14].

All-atom molecular dynamics (MD) simulations have been extensively used as a complementary methodology to experimental approaches, as a means of evaluating the dynamical properties of IDRs and increasing our knowledge of the structural behavior of these regions. Its ability to probe local changes in structural attributes and diverse conformers over time is a valuable source of information to capture significant features of IDRs. However, the application of traditional biomolecular force fields designed for globular regions has been proved limited to obtain accurate molecular dynamics simulations of these regions. As a result, several strategies were developed in recent years to better address some of these issues, making them more reliable for the analysis of IDRs [15], [16], [17].

Since the release of AlphaFold2 [18] and other 3D structure predictors based on deep learning, we are able to finally glimpse a higher volume of structural conformations of IDRs. However, this expansion on the available data brought with it new challenges [19]. AlphaFold2 and other IDR predictors do not always agree on the exact extension and position of the IDRs, and, according to a recent benchmark, no IDR prediction tool presents an accuracy higher than 80% over curated samples [20]. Low (between 50 and 70) or very low values (below 50) of AlphaFold’s per-residue model confidence score (pLDDT) have been suggested as indicative of disorder. Nonetheless, nearly 15% of the residues of predicted human IDRs present a pLDDT higher than 70, with several of those presenting some degree of helical structure. These helical formations are suggested to indicate the capacity of AlphaFold2 to predict conditionally folding regions, which are the conformations adopted by some IDRs upon contact with a ligand or a binding partner. However, this hypothesis requires further investigation, since the number of curated regions that perform conditional folding is still very low, and the selection of negative cases that definitely do not fold upon binding is even harder [21].

All these uncertainties motivated our search for novel ways to evaluate the stability of these helical formations within IDRs. In particular, we focused on polyXY, LCRs composed of a maximum of two different amino acids. In our previous research, we evaluated the general context of polyXYs situated in human IDRs, first exploring their structural context using experimental structural data from the PDB [22], later increasing the structural information available by using AlphaFold2 predictions [23]. We were able to determine that some polyXY types, i.e. EK, ER, DE and AE, show higher helical content. Here, we show that IDRs close to helical structures show accumulation of polyXYs in the boundary close to the helix. To examine this effect, we focused on evaluating the specific subset of annotated polyXYs situated at the ends of IDRs and that are close to helical formations outside the IDR. These polyXYs are particularly interesting because they often extend the helical conformation into the IDR. The composition of these polyXYs was analyzed and finally molecular dynamics simulations were executed to evaluate their helical propensity and stability over time.

## Methods

### Selection and analysis of helical and non-helical neighboring regions

We followed the process executed in our previous work [23] for annotation of human IDRs, polyXYs, and secondary structures using AlphaFold2 predicted regions and the DSSP database of secondary structure assignments. The newest releases of the source databases were used, with MobiDB version: 5.0 - Release: 2022_07 [24] and AlphaFold2 version 4 [25], [26]. DSSP version 3.0.0, was also used, including a distinctive characterization of polyprolines (PPII helices) [27]. The cut-offs applied in the previous work were maintained, with polyXYs composed of a minimum of 6 residues, overlapping with the containing IDR in at least 4 residues or 50% of the polyXY region. IDRs present a minimum size of 20 residues, as set by the prediction methods.

Regular expressions were then applied at both ends of the IDR to ascertain the presence of a helical formation adjacent to the IDR. When located at the N-terminal of the IDR, these helices should contain at least four residues (1 turn) before the IDR and end inside the IDR or at a maximum distance of 3 residues before the IDR. Symmetrical rules were applied to select helical formations at the C-terminal of the IDR (See figure 1-i to iii for a detailed schematic). IDR ends were labeled as “Helical” and “Non - Helical” depending on whether they contained one of these adjacent helices (Supplementary table 1).

**Figure 1.**
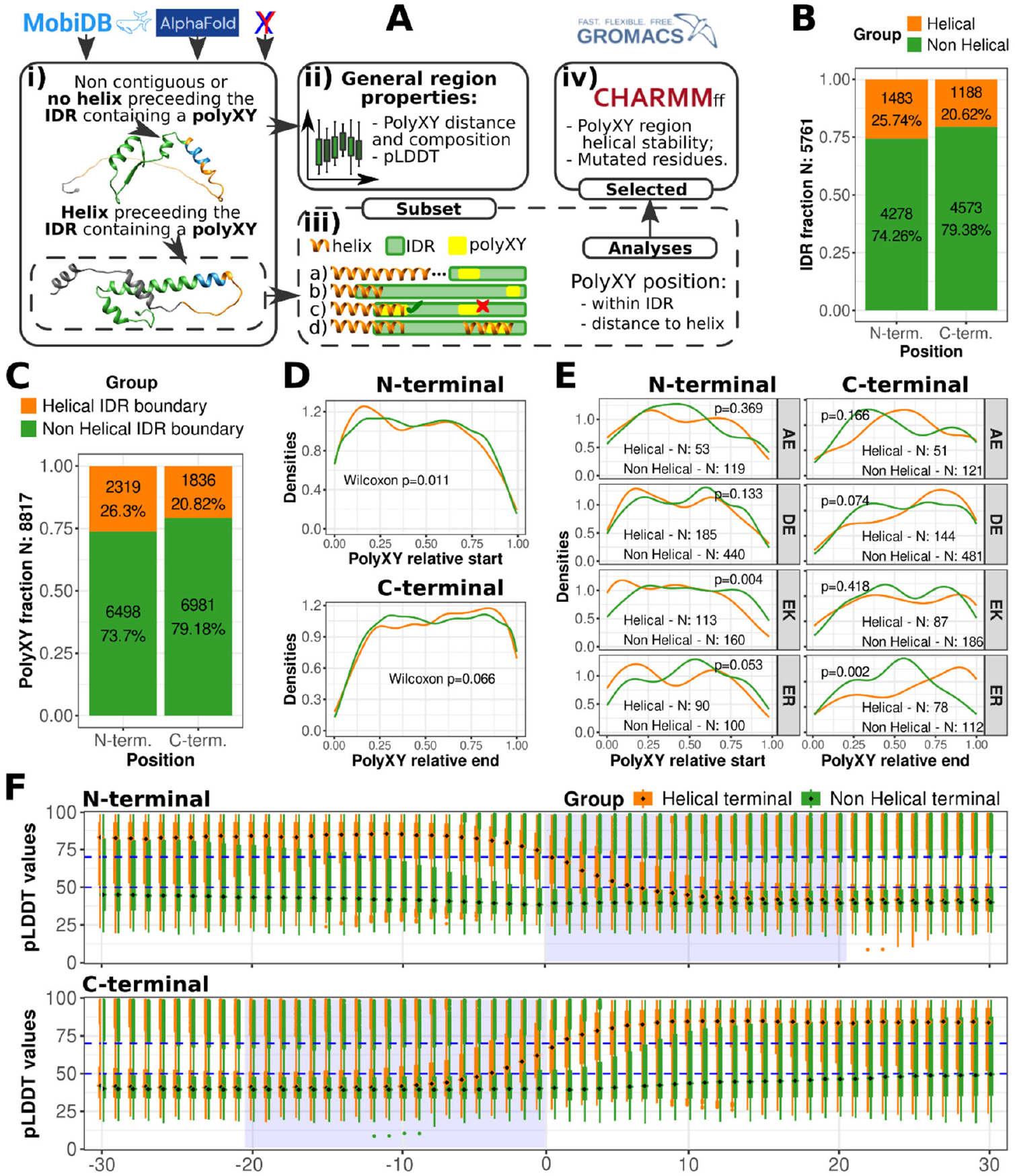
Overview of the neighboring helices of IDRs containing polyXYs. **(A)** A schematic of the process followed to analyze and explore the properties of (i and ii) IDRs containing or not containing an adjacent helix; (iii) polyXYs within IDRs in the context of neighboring helices, considering different IDR, helix and polyXY positions along the region; and (iv) selection and submission of helical polyXYs for MD simulations with the aim of analyzing the helical stability over time. The subclassification in (iii) states for a) helices that end at most three residues before the IDR containing a polyXY begins; b) helices that partially or completely overlap with IDRs, regardless of the polyXY position; c) additional polyXYs within the same IDR were discarded from the second-level analysis, keeping only the first polyXY regardless of whether it overlaps a helix or not; d) even when multiple helices overlap with the same IDR they are kept. **(B)** Counts and percentages of IDRs and **(C)** polyXYs within IDRs according to the presence or absence of a neighboring helix at each IDR termini. **(D)** Distribution of polyXYs globally and **(E)** among the top four polyXYs with highest helical content, considering the relative position within IDR coordinates and the presence or absence of a helix at each IDR termini. **(F)** Average pLDDT values per residue (diamonds) in the vicinity of the IDRs, considering 30 residues before and 30 residues inside the IDRs. The blue shaded box indicates the minimum region covered by all IDRs (20 residues), while the dashed blue horizontal lines delimit regions of intermediate and high confidence, respectively.

### Subsetting polyXYs overlapping IDRs and helices at their termini

As our analysis was focused on the relationship between polyXY within IDRs and neighboring helical structures, we segregated the IDRs containing a helical neighbor for the following analyses. Considering that one IDR can contain several polyXYs, c) we selected the polyXY that was the closest to the target helix, d) independently of whether an overlap between helix and polyXY occurs (Figure 1-iii, Supplementary table 2-3).

Depending on the overlap of the polyXYs with the target helix, we classify them as “high helical overlap” (if they overlap in 6 or more residues), and “low/no helical overlap” (figure 1A-iii). Further analyses were performed considering the proximity between the polyXY center and the IDR or helix ends.

### Additional annotation of biological functions and sequence features

We analyzed the composition of the selected subsets of polyXYs and additional information was collected using the MobiDB database regarding closest protein domains for both IDR ends. For the cases with a helix at the IDR’s N-terminal, the distance between the center of the polyXY and the nearest domain N-terminal to the IDR were collected, with distance 0 if the domain overlapped the polyXY center. The same parameters were mirrored for the C-terminal analyses.

### Selecting and formatting targets for MD simulations

Despite recent advances in computational power and capacity, the execution of all atom MD simulations remains computationally expensive. Therefore, the selection of targets for our analysis was inevitable. First, we selected polyXYs at helical N-terminal IDR ends and considered those with high helical coverage. Of these, we took the 55.6% that are located 20 residues or less from the helix end. We filtered them further considering those with an average pLDDT value greater than 70.

From the resulting set, we selected two samples from each of the three polyXY types that overlap helices most frequently. Before proceeding, a visual inspection of the fragment structure was performed to confirm the helical structure of the region. A segment of 100 residues surrounding the polyXY was sliced from the AlphaFold2 prediction and submitted to simulation. Our decision about using a fragment of the sequence was taken to reduce the execution times, which increase with the size of the molecule under study, and to reduce the risk of artifacts being generated by force fields designed for IDPs when applied to structures that also contain stable globular regions [28]. In all three selected polyXY types that commonly overlap helices, at least one of X or Y is a charged residue. To explore if charge is enough to keep this helical conformation, we experimented with mutations of the polyXYs to the counterpart charged residues. An AlphaFold2 prediction for the mutated protein was generated, then also sliced to 100 residues surrounding the polyXY and submitted to MD simulations with the same conditions.

### Molecular Dynamics simulations and analyses

Gromacs 2022.5 with the all-atoms force field CHARMM36IDPS [29] was used for all simulations. CHARMM36IDPS was optimized using the grid-based energy correction map (CMAP) method, using backbone dihedrals collected from the PDB database, differentiating between globular and non-globular regions. For the setup of the system, cubic boxes with distance between the solute and the box wall of 2.0 nm with periodic boundary conditions were filled with TIP3P water [30], and sodium chloride concentrations were adjusted to human physiological conditions of 100 mM. Steepest-descent algorithm was used for the energy minimization step, with a maximum of 50,000 steps or the energy difference is less than 1000 kJ mol^-1^ nm^−1^ and integration time step of 1 fs. Particle-mesh Ewald (PME) was used to calculate the long-range electrostatics with a cubic interpolation, with a cut-off radius of 1.0. The modified Berendsen thermostat algorithm was used for temperature coupling at 300K and the Parrinello-Rahman algorithm was used for pressure regulation, both steps executed each 100 ps. Production simulations were executed for 1.2 μs, with the first 100 ns of each simulation discarded to reduce the bias that can be caused at the initial states by the AlphaFold2 predicted structures.

The simulation trajectories were then analyzed using Gromacs dssp function with dssp mode version 3.0.0 for the creation of hydrogen pseudo atoms [27]. The canonical MD simulation metrics root mean squared deviation (RMSD) and surface area analysis (SASA) in the polyXY region were also generated to support the findings observed in the secondary structure analyses of the first MD simulation and its mutations. These metrics present global average information on how distant the proposed structures along the trajectories are and how much of these structures are exposed to the surrounding solvent, respectively. To obtain local measurements, our analysis focused on the target helical region and was calculated over a sliding window of six residues centered on the target residue.

### Additional tools

All data was processed through in-house scripts developed in Python 3.8.10. The package Biopython was used to manipulate FASTA files and extract DSSP annotations [31]. All plots and tables were generated with scripts in R 4.4.0 and ggplot2 3.5.1, with p-values calculated using Mann-Whitney U Test (Wilcoxon Rank Sum Test). Protein molecular structures were generated with Chimera 1.15 [32] and MD simulation analyses were performed over Gromacs native reports, with plots generated with Python scripts.

## Results

### IDRs neighboring helical structures show accumulation of polyXYs close to the helices

Our total set of human IDRs containing polyXYs (5761 - 23.31% of all human IDRs) was divided into two groups, containing and not containing helical structures adjacent to the IDR (see Methods for details). As proteins are synthesized from N- to C-terminal, it is feasible that secondary structures N-terminal to the IDR are already formed when the IDR starts to be synthesized, even within the ribosomal tunnel [33]. This could impose differences in the structural propensities of each IDR boundary, for example facilitating the formation of structures at the N-terminal. For this reason, we contemplated separately the N- and C-terminals (Supplementary table S1). Consistent with this hypothesis, we observed that a slightly higher percentage of IDRs containing polyXYs present helices at their N-terminal (25.74%) compared to their C-terminal (20.62%) (Figure 1B).

We then evaluated the polyXY content of these IDRs. Considering IDRs preceded by helices at their N-terminal, 2319 polyXYs were selected, with an average of 1.56 polyXYs per IDR, while a total of 1836 polyXYs with a similar average of 1.55 polyXYs per IDR can be observed for IDRs with helical formations at the C-terminal (Figure 1C).

To further explore the relationship between polyXYs and helicity, we investigated whether IDRs preceded or followed by a helix presented an accumulation of polyXYs closer to the IDR end and the corresponding adjacent helix, compared to the cases where no helix is present in the immediate vicinity of the disordered region.

We show that this is the case: an accumulation of polyXYs within the group of IDRs neighboring helices can be observed compared to the cases where no helix is present, with a significant p-value = 0.011 for the N-terminal and a non-significant p-value = 0.066 for the C-terminal, according to the Wilcoxon rank test, with a threshold of p<0.05 (Figure 1D). The top four polyXYs that presented higher counts of helical content amongst polyXYs within IDRs according to our previous work [23] were also evaluated. For the cases with an IDR with a helical N-terminal, polyAE and polyDE present a lower accumulation closer to the start of the IDR, while EK and ER present a higher accumulation, with the first presenting significant p-value < 0.01 (Figure 1E). This trend is attenuated for the C-terminal samples, except for polyER, with significant p-value < 0.01, and polyDE with a non-significant accumulation to the end of the IDR, indicating a more scattered distribution of polyXY types at the C-terminal.

Considering the presence of a helical boundary, one would expect that AlphaFold confidence levels would be higher at the IDR ends, expanding at least marginally within the IDR, compared to samples where no boundary helix is present. The pLDDT distributions were accessed for both termini (Figure 1F), and these expectations were confirmed, with an average pLDDT above 75 for residues before the IDR border and between 50 and 75 for the 6 first residues at the N-terminal and 4 residues at the C-terminal of the IDRs.

### Charged and helical prone polyXYs show a higher helical propensity within helical IDRs ends

The following analyses were directed to the polyXYs within IDRs with helical ends. Considering that one IDR can contain several polyXYs, just the first and last polyXY, closer to the IDR ends, were selected (Supplementary Tables S2-S3). We analyzed their residue composition and their position relative to the helix advancing into the IDR.

We evaluated 1435 polyXYs at the N-terminal, of which 225 (16%) were covered by 6 or more helical residues (high helical coverage; Supplementary Table S2). A similar distribution of polyXYs regarding their helical coverage could be observed at the C-terminal, with a total of 1153 polyXYs, of which 160 (14%) had high helical coverage (Supplementary Table S3). We observed that while polyXYs with helical coverage are significantly closer to the surrounding helix at the N-terminal, this is not the case at the C-terminal (Figure 2). We interpret this result as indicating a stronger association of polyXYs with helical structures placed at the N-terminal of the IDR than those at the C-terminal; such N-terminal helices could then be extended into the IDR with the assistance of the polyXY.

**Figure 2.**
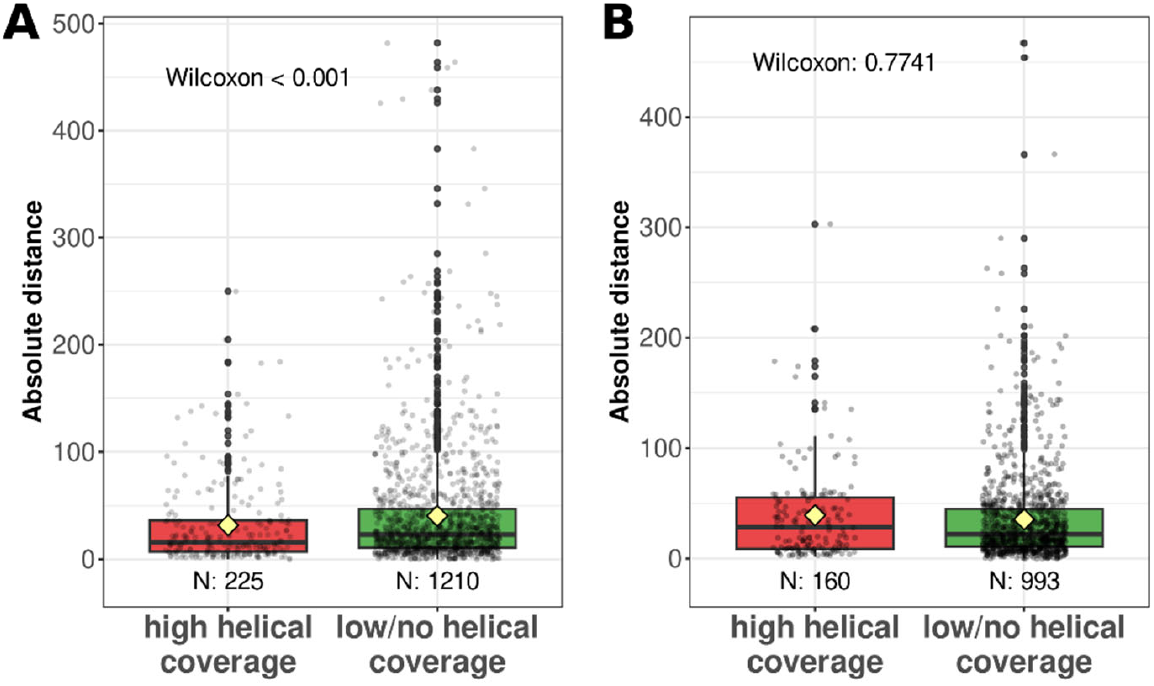
Distances between polyXYs and helices surrounding IDRs. Distances from the center of the polyXY to **(A)** the end of the helix at the N-terminal of the IDR, and **(B)** to the start of the helix at the C-terminal of the IDR, considering just IDRs containing an adjacent helix. Total of polyXYs for each group is shown. Two-sided Wilcoxon statistical test was applied, considering as the null hypothesis that there are differences between the two groups of polyXYs with “high helical coverage” and “low/no helical coverage”.

We next assessed the residue composition of these polyXYs close to helical IDR ends (Figure 3). The most frequent polyXY types were similar at the N- and at the C-terminals. For low/no helical coverage, polyDE and polyAP are the most frequent types, followed by polyXY types containing serine, glycine and proline. For high helical coverage, glutamic containing polyXY are the most frequent: polyER, polyEQ, polyEK and polyAE.

**Figure 3.**
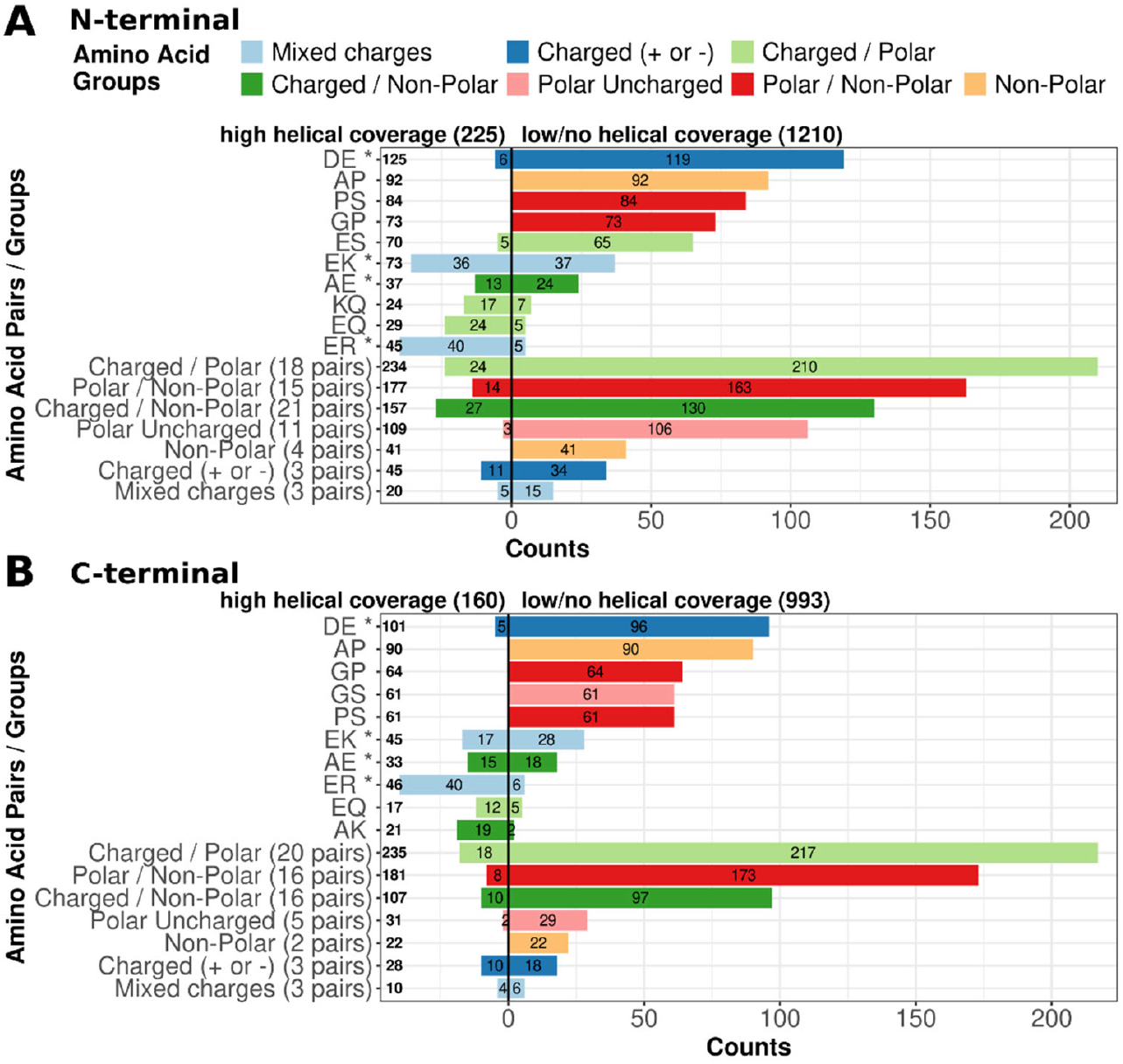
Composition of polyXYs within helical IDR ends. Counts of polyXYs grouped as “high helical coverage” on the left and “low/no helical coverage” on the right. The top five polyXYs showing the highest counts in each group are detailed, while the remaining polyXY types were further grouped by physicochemical composition of the residue pair, indicated by color. The top four polyXYs containing the highest overall helical coverage according to the previous study [23] were highlighted with an asterisk. **(A)** Helical N-terminal ends, **(B)** helical C-terminal ends.

In the next section, to complement our findings obtained from sequence analysis and structural data, we apply MD simulations to some examples selected among the polyXY types most prominently observed as associated to IDR helical ends, (i) to see if we can find dynamic information for some representative polyXY structures and (ii) to test the resilience of these structures to conservative mutations.

### MD simulations of TERA polyER and variants

To analyze the dynamics and stability of an example of polyER situated at a helical N-terminal IDR, we focused on TERA, which, according to an electron microscopy solved structure, presents a transient helical coverage along the polyEK (PDB:7bpa; Figure 4A)[34]. TERA stands for transitional endoplasmic reticulum ATPase (UniProtAC: P55072; also known as p97 and VCP). This protein is a hexameric AAA+ ATPase largely investigated due to its participation in several biological functions, including endoplasmic reticulum associated protein degradation and ubiquitin regulation. Due to its relevance to pathways related to cellular stress responses, some of its mutations are associated with diseases such as Amyotrophic lateral sclerosis and Paget’s disease of bone; and it is upregulated in certain cancer types [35], [36]. A few residues following the polyER are missing in the solved structure (Figure 4A, inset) suggesting that these adopt a flexible conformation. Examination of a total of 111 structures of TERA monomers deposited in the PDB shows 38 cases where the polyER is also completely helical, 13 cases where it is partially helical and 30 cases where it is flexible, suggesting that the polyER might also be flexible.

**Figure 4.**
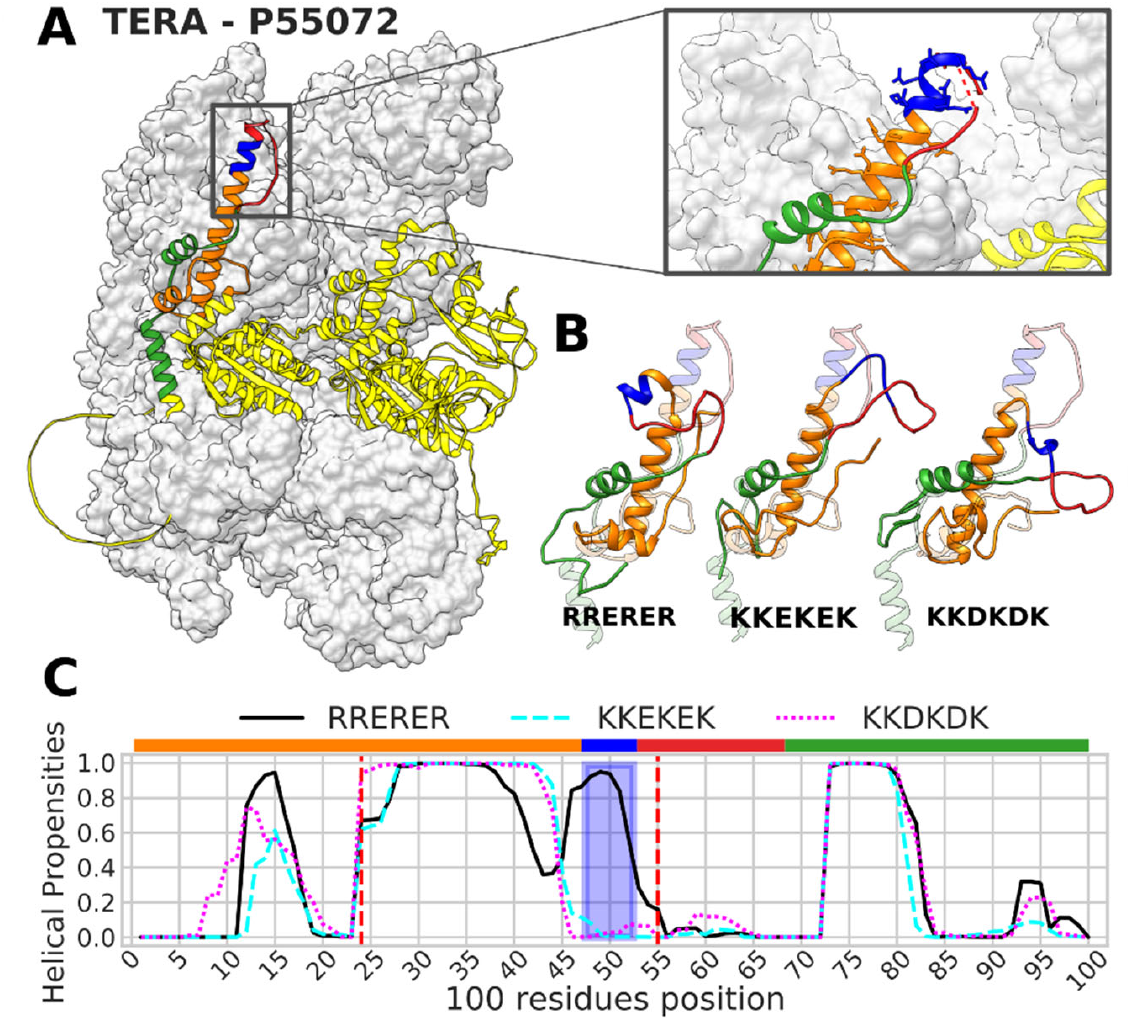
TERA structure and MD simulation conformations after 1.2 μs for original and mutated polyER. **(A)** Electron microscopy of the resolved structure of the TERA hexamer (PDB:7bpa)[34], with AlphaFold2 predicted structure for TERA superimposed on chain D of the solved structure. Colors indicate: polyER (blue), IDR (red), and the 100-residue region included in the MD simulations (Orange and green for N-and C-terminal, respectively). The inset shows the original solved structure: the discontinuous line indicates residues missing, suggesting that a few residues C-terminal to the polyER might adopt a flexible conformation. **(B)** The structure of the 100-residue region surrounding the polyER was predicted with AlphaFold2 for the original sequence and for two mutated polyXYs as indicated (faded colors) and MD simulations were applied for 1.2μs (final structures in stronger colors). **(C)** Helical propensities during the MD simulations. The blue box delimits the polyER region and the red dashed vertical lines indicate the position where the helix originally started and ended in the AlphaFold2 prediction. The first 100 ns were discarded to remove a possible bias from the structure originally predicted by Alphafold2.

To further explore the stability of the predicted helix and the conformational variability of the segment, we ran 1.2 μs long MD simulations for a region of 100 residues surrounding the original polyER, RRERER, and to two mutated polyXYs: KKEKEK, replacing the Rs by similarly positively charged Ks; and KKDKDK, replacing both residue types by similarly charged residues. PolyEK is, like polyER, a frequent polyXY with high helical coverage (Figure 3), while polyDK is less common. The resulting structures of the MD simulation indicate that the original polyER region, even when not exposed to the hexameric structure and ligands, still presents traces of a helical formation, while the two mutated structures completely lost the helical structure up to four residues before the polyXY (Figure 4B).

Using the detailed propensities over the MD trajectory, we can analyze the behavior of specific residues in the helical segment surrounding the polyER (Supplementary figure S1). Generally, some instability of the helical region of six residues preceding the polyER can be appreciated starting at 350 ns, with most residues keeping some degree of helical propensity (Supplementary figure S1A). Something similar can be observed for the first four residues of the polyER but the last two residues and the next two lose helical propensity already at 50 ns (Supplementary figure S1B). The MS trajectories for the sequence mutated to polyEK present a different scenario (Supplementary figure S2A). The three residues preceding the polyEK lose helical structure at 200 ns (Supplementary figure S2A), and the polyEK and following residues at 50 ns (Supplementary figure S2B). A similar scenario is observed in the MS trajectories for the polyER sequence mutated to polyDK, where the three residues preceding the polyDK, the LCR itself and the next two residues lose the helical structure at 150 ns (Supplementary figure S3A-B).

RMSD and SASA, two canonical MD simulations metrics, were also generated to allow the comparison between the polyXY regions and the adjacent helix, which presented differences in helical propensities among the three simulations. While there are dramatic differences in helical propensity between the original sequence and those of the mutated polyXY (Supplementary figure S4A), both RMSD and SASA values per residue are very similar. In all three simulations, we can observe that the RMSD values and standard deviation are higher in the polyXY than in the preceding helix but lower than in the following IDR residues (Supplementary figure S4B), while the SASA values are the highest in the polyXY (Supplementary figure S4C). We interpret this result as indicating that the changes in sequence affect specifically the helical structural propensity of the sequence, without altering primary properties that describe the dynamics of the sequence in the simulation.

These results suggest that the short polyER contributes to the stability of the preceding helical residues, and replacement by other residues, even with similar physicochemical properties, does not favor the same structure. The partial loss of structure before the polyER observed in the simulation of the original sequence can be the result of the absence of interacting partners, which are present in all experiments performed with this hexameric protein. While the polyER does not interact directly with the neighbor chain, the two glutamic acid residues in the preceding helical region interacts, making the helical formation at this region essential for the stability of the protein[34]. Taken together, the stability noticed in the original sequence and the AlphFold2 prediction reinforce the traits of a region with inherent flexibility and helical propensities.

### MD simulations of other polyXYs with helical propensities

To expand the results observed with TERA, we selected examples of other frequent polyXYs with high helical coverage situated at helical IDR N-terminals, all of them with a median pLDDT within the polyXY region above 70 (see Methods for details): one additional polyER, two polyEK and two polyEQ (Supplementary table S4), and subjected them to MD simulations. Unlike for TERA, no solved structures are available for the helical region or the corresponding polyXYs and we rely on AlphaFold2 for structural information.

The Heparan-sulfate 6-O-sulfotransferase 3 protein (HS6ST3; UniprotAC: Q8IZP7) is a 6-O-sulfation enzyme that catalyzes the transfer of a sulfate of heparan sulfate. The polyER (RREERR) targeted in our analysis belongs to the C-terminal IDR of the protein, which follows a Sulfotransferase domain (Supplementary table S4). Due to its position within the sequence, the simulation was performed over a fragment of 85 residues. While the AlphaFold2 predicted structure presents a long 46 residue helix spanning almost half the IDR and its N-terminal region, our MD resulting structure shows a shorter helical formation, with the region at the C-terminal of the polyER completely unfolded along with most of the fragment at the N-terminal of the IDR (Figure 5A). Evaluating the helical propensities of the whole simulation, the same scenario can be observed, with a decrease of the helical structure at the end of the polyER, still remaining helical for around 60% of the trajectories (Figure 5B). When evaluating the helical stability of the polyER and the next two residues over time (Supplementary figure S5), we can observe that the end half of the polyER had a reduction of the helical propensities for most of the simulation, with a complete loss between 400 and 550 ns, later recovering its helical structure around 800 ns and remaining helical until the end of the simulation.

**Figure 5.**
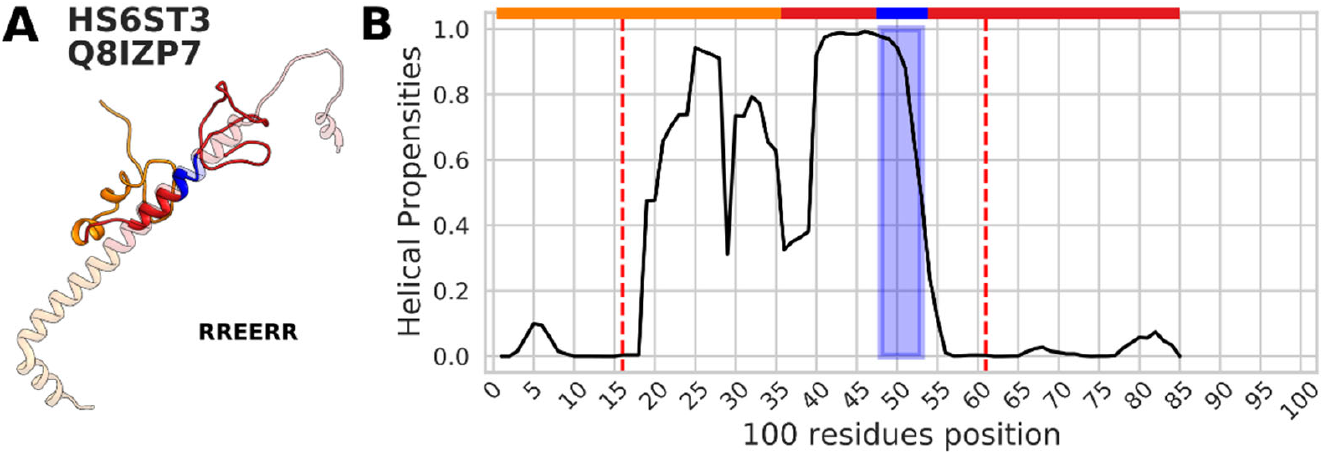
MD simulation of a segment of HS6ST3 containing a polyER region. **(A)** Superimposition of the original AlphaFold2 prediction for a fragment of 85 residues surrounding the polyER region (faded colors) with the structure resulting from the MD simulations after 1.2 μs (shown in stronger colors). Colors indicate the polyER region (blue), the remainder of the IDR (red) and the region reaching to the 50 residues N-terminal to the polyER (orange). The segment includes the end of the protein sequence. **(B)** Helical propensities for the duration of the MD simulation (first 100 ns were discarded). The blue box delimits the polyXY region and the red dashed vertical lines indicate the position where the helix originally started and ended in the AlphaFold2 prediction. The color bar above delimits the fragment regions as described in (A).

Our first polyEK sample (KKEKKEKK) is from the RNA polymerase-associated protein RTF1 homolog (UniprotAC: Q92541). The 100-residues region surrounding the polyEK has a long IDR, which covers 59 residues including the polyEK and the whole C-terminal of the segment. The resulting structure of the MD simulation presents helical coverage only at the two last residues of the polyEK (Figure 6A). The residues in the polyEK present values of helical propensity above 50% (Figure 6B). However, the MD trajectories show that at 1 μs only the last two residues remained helical (Supplementary figure S6B). All residues from the adjacent region at the N-terminal of the sequence also lost completely the helical coverage at 1 μs (Supplementary figure S6A). An EK rich region that follows the polyEK has high helical propensity (KKQEEEQE) and remained mostly helical throughout the whole simulation (Supplementary figure S6C). This region might be acting as an extension of the considered polyEK region.

**Figure 6.**
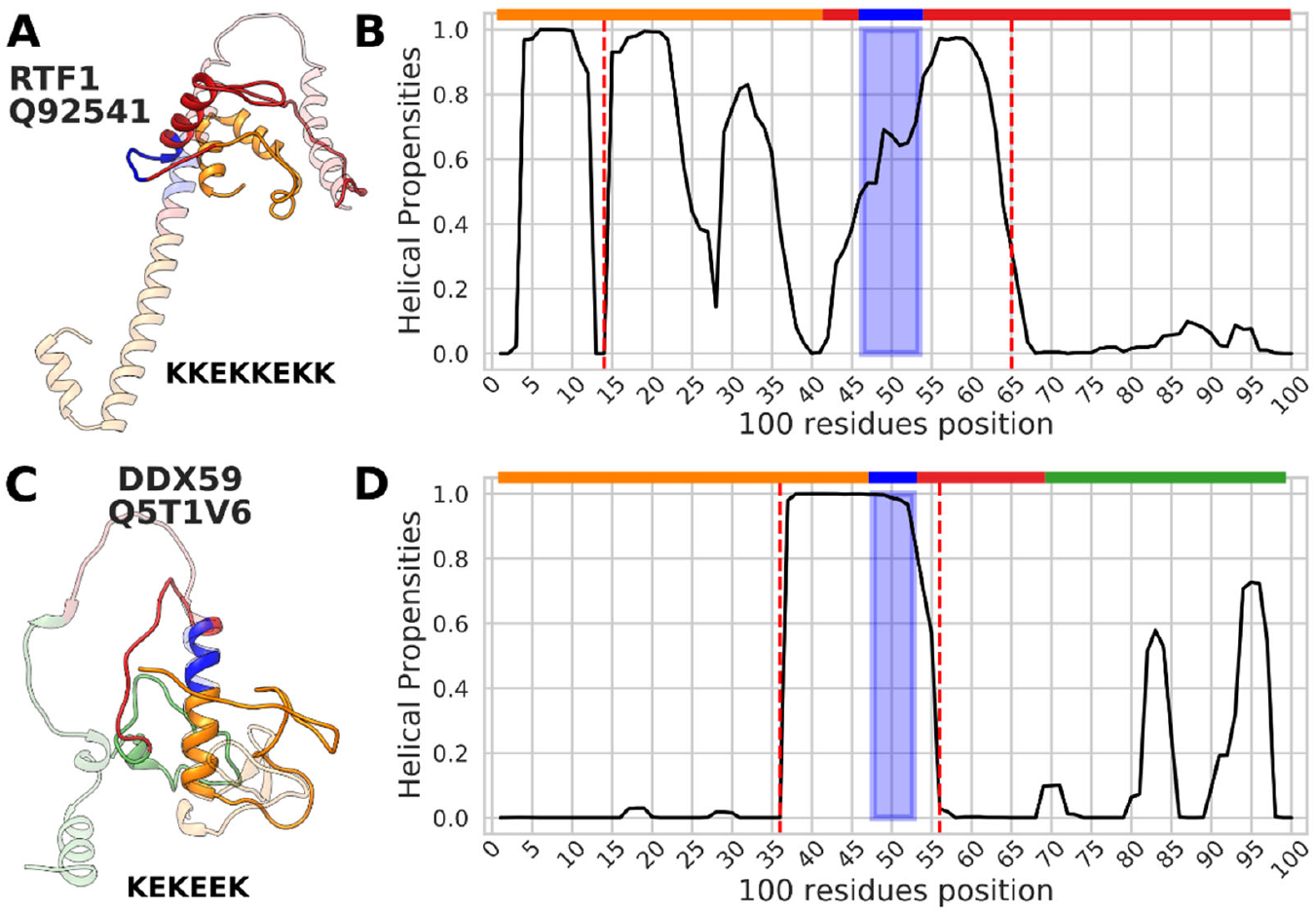
MD simulations for two segments containing a polyEK region. **(A)** Superimposition of the original AlphaFold2 prediction for a fragment of 100 residues of RTF1 surrounding the polyEK region (faded colors) with the structure resulting from the MD simulations after 1.2 μs (shown in stronger colors). Colors indicate the polyEK region (blue), the remainder of the IDR (red) and the region reaching the 50 residues N-terminal to the polyEK, which also contains the preceding helix (orange). **(B)** Helical propensities for the duration of the MD simulation of RTF1. The first 100 ns were discarded to remove a possible bias from the structure originally predicted by AlphaFold2. The blue box delimits the polyXY region and the red dashed vertical lines indicate the position where the helix originally started and ended in the AlphaFold2 prediction. The color bar above delimits the fragment regions as described in (A). **(C)** Superimposition of the original AlphaFold2 prediction for a fragment of 100 residues of DDX59 surrounding the polyEK region (faded colors) with the structure resulting from the MD simulations after 1.2 μs (shown in stronger colors). Colors indicate the polyEK region (blue), the remainder of the IDR (red), the region reaching to 50 residues to the N-terminal of the polyEK center, which also contains the preceding helix (orange) and the region reaching to the 50 residues to the C-terminal of the polyEK center (green). **(D)** Helical propensities for the duration of the MD simulation of DDX59. The first 100 ns were discarded to remove a possible bias from the structure originally predicted by Alphafold2. The blue box delimits the polyXY region and the red dashed vertical lines indicate the position where the helix originally started and ended in the AlphaFold2 prediction. The color bar above delimits the fragment regions as described in (C).

The second analyzed polyEK was from DDX59 (UniprotAC: Q5T1V6), a probable ATP-dependent RNA helicase. The center of the targeted polyXY region (KEKEEK) is five residues distant from the end of the N-terminal helix, with a Zinc finger domain ending eight residues before the start of the polyEK (Supplementary table S4). The polyEK region presented a remarkably stable helical composition throughout the whole 1.2 μs, with some loss of structure in the last residue of the polyXY and in the following three residues (Figure 6C-D). This loss occurs already at 100 ns of the simulation (Supplementary figure S6D).

The first selected polyEQ (QQEQEQ) was from PIEZO1 (UniprotAC: Q92508), a pore-forming subunit of the cation Piezo channel. After 1.2 μs of MD simulation, the helical segment which includes the polyEQ and a few surrounding residues, remains stable (Figure 7A). The helical propensities are high in the first section of the polyEQ, with a reduction to 70% at the end section and lower after the polyEQ (Figure 7B). We can observe the loss of helical structure at the last three residues of the polyEQ between 400 ns and 550 ns, with a full recovery at the end of the trajectory (Supplementary figure S7A). Within the same timeframe, the five residues following the polyEQ also lost the helical structure, gradually recovering afterwards (Supplementary figure S7B).

**Figure 7.**
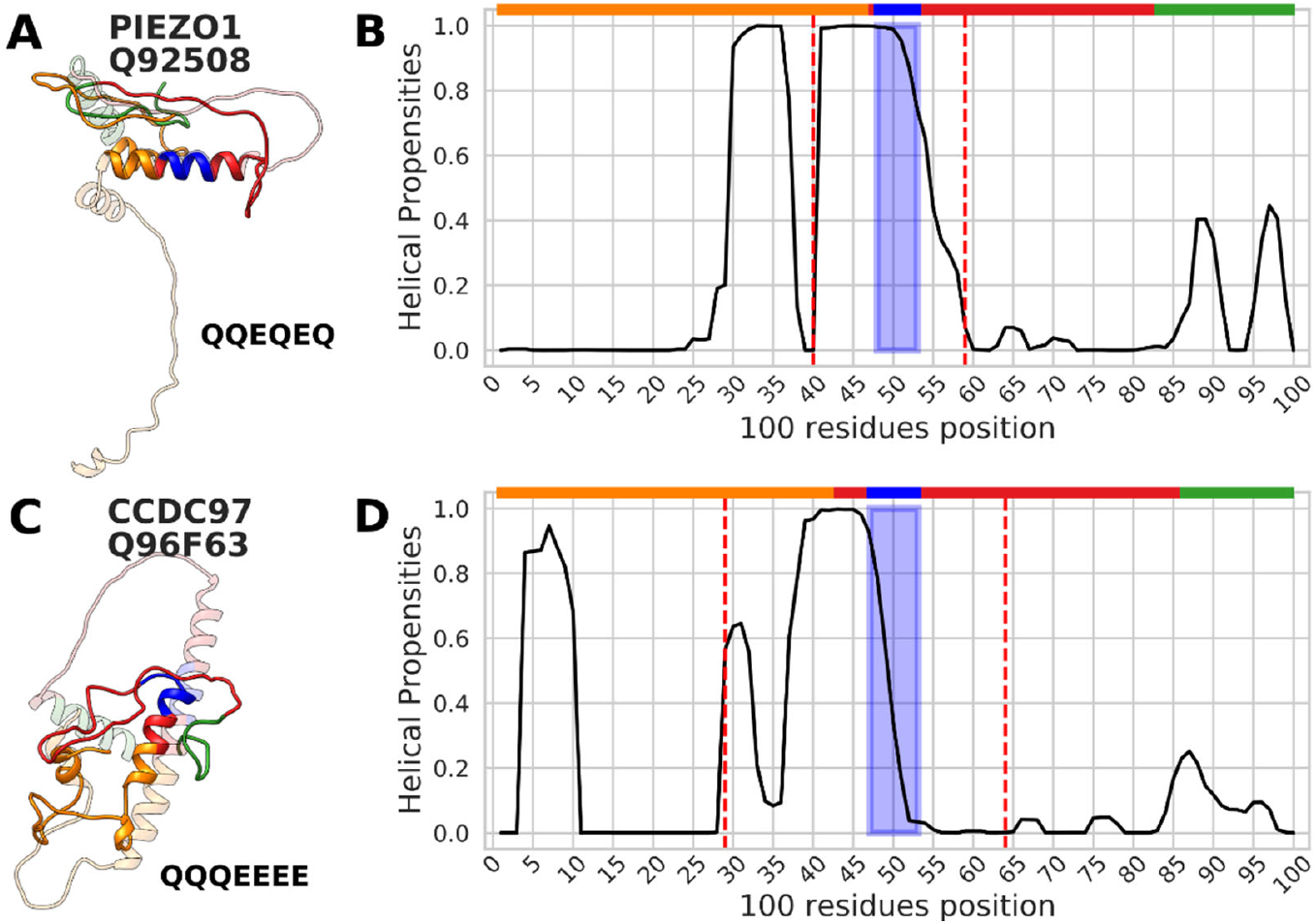
MD simulations for two segments containing a polyEQ region. **(A)** Superimposition of the original AlphaFold2 prediction for a fragment of 100 residues of PIEZO1 surrounding the polyEQ region (faded colors) with the structure resulting from the MD simulations after 1.2 μs (shown in stronger colors). Colors indicate the polyEQ region (blue), the remainder of the IDR (red), the region reaching to 50 residues to the N-terminal of the polyEQ center, which also contains the preceding helix (orange) and the region reaching to 50 residues to the C-terminal of the polyEQ center (green). **(B)** Helical propensities for the duration of the MD simulation of PIEZO1. The first 100 ns were discarded to remove a possible bias from the structure originally predicted by Alphafold2. The blue box delimits the polyXY region and the red dashed vertical lines indicate the position where the helix originally started and ended in the AlphaFold2 prediction. The color bar above delimits the fragment regions as described in (A). **(C)** Superimposition of the original AlphaFold2 prediction for the 100 residues of CCDC97 surrounding the polyEQ region (faded colors) with the structure resulting from the MD simulations after 1.2 μs (shown in stronger colors). Colors indicate the polyEQ region (blue), the remainder of the IDR (red), the region reaching 50 residues to the N-terminal of the polyEQ center, which also contains the preceding helix (orange) and the region reaching 50 residues to the C-terminal of the polyXY center (green). **(D)** Helical propensities for the duration of the MD simulation of CCDC97. The first 100 ns were discarded to remove a possible bias from the structure originally predicted by AlphaFold2. The blue box delimits the polyXY region and the red dashed vertical lines indicate the position where the helix originally started and ended in the AlphaFold2 prediction. The color bar above delimits the fragment regions as described in (C).

Our last sample, a second polyEQ (QQQEEEE), is from CCDC97 (UniprotAC: Q96F63). Here, helical stability can be observed in the glutamine portion of the polyEQ, which kept a propensity of more than 40% throughout the whole simulation along with the region before the polyEQ (Figure 7C-D). Interestingly, among all the segments chosen for simulations, this is the only one that combines two homorepeats and also presented the highest loss rate of helical content throughout the polyXY region. Additional simulations would be required to investigate the relationship between the order of the residues within a polyXY and their helical stability. The glutamic acid segment of the polyEQ lost most of its initial conformation in the simulation process already at 200 ns with a gradual recovery at the end (Supplementary figure S7C). The polyE following the polyEQ predicted by AlphaFold2 as helical, lost completely the conformation at early stages of the simulation (Supplementary figure S7D).

In summary, most of the polyXYs studied in the selected fragments presented helical propensity values above 50% at the end of the 1.2 μs simulation, with a tendency to lose structure at the C-terminal (away from the N-terminally situated adjacent helix). These findings support the predictions presented by AlphaFold2 in these particular regions. The predictions for the helix preceding the polyXY were less stable, except in the case of DDX59 (Figure 6C-D) and PIEZO1 (Figure 7A-B).

## Discussion

In previous work, short LCRs composed of one or two amino acid types, denominated polyX and polyXYs, respectively, were characterized, exploring their position within protein sequences and IDRs, residue preferences, categories and protein biological functions [37], [38]. Their structural properties when within IDRs were assessed for the human proteome, initially through homology to PDB structures, with some residues presenting helical propensities, including E and K, alone or associated, or helical avoidance, including polyEP and polyGX [22]. AlphaFold2 predictions increased the pool of structured candidates, not only confirming our previous findings for helical prone structures, but also adding some new LCRs to this list, such as polyQ and polyER [23].

The importance of studying helical propensities, also called residual structures, within IDRs has been explored. The capacity to rapidly adopt a specific secondary structure upon-binding, a folding pathway called conformational selection, would increase the affinity of interactions between some disordered and partner regions [39]. There are discussions on how these regions benefit the formed complexes, whether enhancing the complex association rates and molecular recognition [40] or by reducing dissociation [41]. Nonetheless, these propensities are important to optimize IDR interactions. Short LCRs are not the only regions capable of exhibiting such propensities, but they may allow them to occur. Shorter LCRs are often inserted in longer and less complex LCR stretches [42]. A better understanding of their local structural behavior can provide invaluable insights into how they can support the function of the longer biased regions. Despite the advancements on the qualification and quantification of these short LCRs within IDRs, the understanding of their specific local structural function still evades us.

AlphaFold2 presented an extensive advance on what we now know about the relationship between IDRs, LCRs and secondary structure. Its predictions are, however, known for sometimes not being realistic and present an overrepresentation of helical formations. These characteristics can be noticed especially in IDR regions, where low pLDDT scores indicate a lack of confidence in the adopted structures [19] and in the large increase of helical structures within IDRs [43], [44], explained as the result of the training over a large set of protein-bound structures, thus not representing the unbound structure in solution [21], [45], [46].

These new challenges presented by AlphaFold2 came as an opportunity to evaluate the behavior of polyXYs within IDRs and to assess their local function. Here, we propose a specific function for a specific group of polyXYs: The extension and maintenance of helical structures formed outside of the IDR region. Targeting this specific scenario would also bring some insight on the behavior of a transition region, often neglected by studies targeting only IDRs or globular regions separately. The amino acid composition of IDRs’ neighboring regions was recently explored, presenting characteristics that suggest a different nature compared to both disordered and ordered regions, evolved and selected to support the structural transitions allowing order to become disordered [47]. In this work, we explored the inverse perspective: how ordered regions can bring order to disordered regions and why this transition is meaningful.

### The categorization of a subgroup of polyXYs within IDRs

To address our hypothesis, we first detected fitting candidates for our analysis, which naturally fell in a specific group of polyXYs in the vicinity of helices that span the ends of the IDR. This subset of polyXYs presents a similar composition at both IDR ends yet showing higher consistency at the N-terminal samples, an expected result of the natural direction of the synthesis of proteins in the ribosome [33]. AlphaFold2 pLDDTs within those regions also present an expected behavior, considering that those are transitioning regions between order and disorder, showing higher values gradually fading when helices are present at the IDR ends, compared to cases where helices are not present.

Further analyzing the structural composition of the two groups in the target LCRs, with “high helical coverage” and “low/no helical coverage”, we explored their composition, searching for distinctions between this subset of polyXYs compared to the global scenario observed in our previous work. The same polyXY types were detected in this subgroup, however the preferences in helical composition differed, with polyDE mainly avoiding helices, even when they are present at the IDR ends, polyEK presenting mixed preferences, which indicate that the surrounding sequence plays an important role in their distinction; and polyER being overwhelmingly covered by helices when a neighbor helix is present, reaching more than 95% coverage when positioned within 50 residues or 33% of the IDR ends when a helix is present. Different polyXYs show a higher relevance on this subgroup, compared to the whole set of polyXYs covered by helices, with polyAE showing the same versatility as polyEK, and both polyKQ and polyEQ also show a higher preference for helical coverage.

We additionally show that some residues have different frequencies within polyXYs with helical coverage in the vicinity of neighboring IDRs; E, K, R, A and Q showing preference; and D, P, S G avoiding these structures. This proposition is not new, with some of the involved amino acid residues being related to helical propensity in polyX [48], [49], however, here we show what types of polyXY are common or avoided in the specific scenario where helices are present in the vicinity of IDRs according to AlphaFold2 predictions. This characterization may not only be important to define this targeting group but can also support future sub-categorizations over other types of polyXYs within IDRs with helical propensities.

Interestingly, we found significantly lower distances of polyXY regions covered by neighboring helices at helical N-terminal IDRs ends but not at C-terminal IDR ends. Again, we take this as deriving from the direction of protein synthesis [33]: it might be easier to expand an already folded helix into an IDR as it is being synthesized, than propagating a helix into an already completed and flexible IDR.

### Reinforcing the helical preferences predicted by AlphaFold2 and the role of polyXYs in local helical stability

The choice of executing MD simulations to explore the proposed AlphaFold2 helical predictions in IDR regions came naturally, as a technique increasingly used to explore the conformational variety of the structural behavior of IDRs [50]. Our findings with the native TERA protein and with its mutated polyXY versions suggest that its polyER formation at the end of the N-terminal helix acts as a stabilizer, ensuring that the previous helix, essential for interaction, remains at least partially stable in our simulations. The mutated versions of TERA’s polyER (polyEK and polyDK) indicated that the presence of a polyER in the region is absolutely essential for the adoption and maintenance of helical structure at the polyXY and at the residues immediately adjacent to the IDR, residues which directly present intermolecular interactions with the hexamer’s partner protein.

To further explore the helical stability not only on the highly helical polyER, but also of some other polyXY types with helical propensities, we performed simulations for five other polyXY containing segments: one polyER, two polyEKs and two polyEQs. Each of the MD simulations presents distinct characteristics that require additional and focused exploration and understanding. Nonetheless, all of them showed a common aspect: they reinforced, at least partially, the helical preference of the region as predicted by AlphaFold2 and, more importantly to our analysis, they indicate that in all scenarios the helical conformation is sustained in the polyXY region.

Almost all samples showed a degradation of the helical structure towards the end of the polyXY, which may suggest that these short LCRs facilitate the extension of the secondary structure into the IDR but not beyond the LCR. Only the simulation of RTF1 (polyEK) showed an increasing gradient of helical propensity towards the end of the polyXY region. The presence of a stable EK-rich region (Figure 6A-B and Supplementary figure S6C) following the polyEK suggests that the definition of the polyEK region might have been too restrictive in this case.

### Limitations and future perspectives

Despite showing an interesting perspective of local structural function of polyXYs within IDRs in the vicinity of helical structures, our work has some limitations.

Firstly, the number of polyXYs with these characteristics was limited by the specificity of the chosen scenario and definitions. This limitation eased the selection of suitable targets for MD simulation, but also limited our ability to further explore the amino acid composition of the surrounding region in a search for additional evidence on why these regions are able to support helical structures and the interplay between the polyXY and its neighboring residues.

In addition, the number of suitable cases can profit enormously from the addition of polyXYs with the same characteristics in proteins from other eukaryotic species without homology to human proteins. The increase of cases and analysis of the surrounding amino acid composition could also allow strengthening the findings from the MD simulations, which is not possible at the moment with our results, due to the diverse scenarios presented by the targeted samples. With additional cases we could then explore their domain context, binding regions, and charge composition, which may shed light on an additional categorization underlying the types of polyXYs analyzed. The search for longer LCRs in which these polyXYs are embedded, considering charged compositional bias and other different LCR compositions, could also further support this characterization.

Secondly, our approach was strongly affected by its scalability. The MD simulations approach, despite being strongly informative, remains extremely restrictive. Our choice to limit the simulation to a fraction of the protein reduced the simulation execution times and ensured successful executions, however limiting the understanding of the behavior of the whole protein. These fragments are impacted by long range interactions [51], which cannot be accounted for in our analysis. Advancements in computational power and the association of different techniques along with ways to reduce MD simulation execution times [52], [53] and the increasing development of force-fields better suited to proteins with mixed ordered and disordered regions [28], [54] may allow these limitations to be overcome in the future. The association of MD simulations with machine learning models can also improve the quality of the results, providing additional validation to the results obtained [55].

Additionally, new avenues can be explored to answer questions raised by this work. Some examples are further analyses of polyEK and polyAE’s dual preferences for helical and flexible content (Figure 3), and the analysis of flexible polyXYs in close vicinity to the external helix end which may act as barriers for the helical extension. Indeed, a systematic study of the helical stability of all polyXY cases with helical content predicted by AlphaFold2 would be desirable. but it is currently beyond our computational capabilities. It is our understanding that our findings using the combination of computational analysis and MD simulations pave the way for a better understanding of the relationship between short LCRs and structural propensity within IDRs. We have presented approaches that should translate future advances in protein structure resolution and prediction and more powerful and specialized MD simulations into further insights about the structural dynamics of flexible regions of proteins.

## Author contributions

Conceptualization, F.S. and M.A.A.N.; methodology for MD simulations, L.A.B. and M.G.K.; global methodology, M.G.K. and M.A.A.N.; software, M.G.K.; validation, L.A.B., M.G.K., F.S. and M.A.A.N.; formal analysis, M.A.A.N.; investigation, M.G.K.; resources, F.S. and M.A.A.N.; data curation, F.S. and M.A.A.N.; writing—original draft preparation, M.G.K. and M.A.A.N.; writing—review and editing, F.S., L.A.B. and M.A.A.N.; visualization, M.G.K.; supervision, F.S. and M.A.A.N.; project administration, M.A.A.N.; funding acquisition, F.S. and M.A.A.N. All authors have read and agreed to the published version of the manuscript.

## Data availability

Data will be made available upon request.

## Acknowledgments

The authors gratefully acknowledge the computing time granted on the supercomputer MOGON 2 at Johannes Gutenberg University Mainz (hpc.uni-mainz.de) for AlphaFold2 predictions and MD simulations.

