## Supplementary Materials for "Assessing the helical stability of polyXYs at the boundaries of Intrinsically Disordered Regions with MD simulations"

### Supplementary Figures

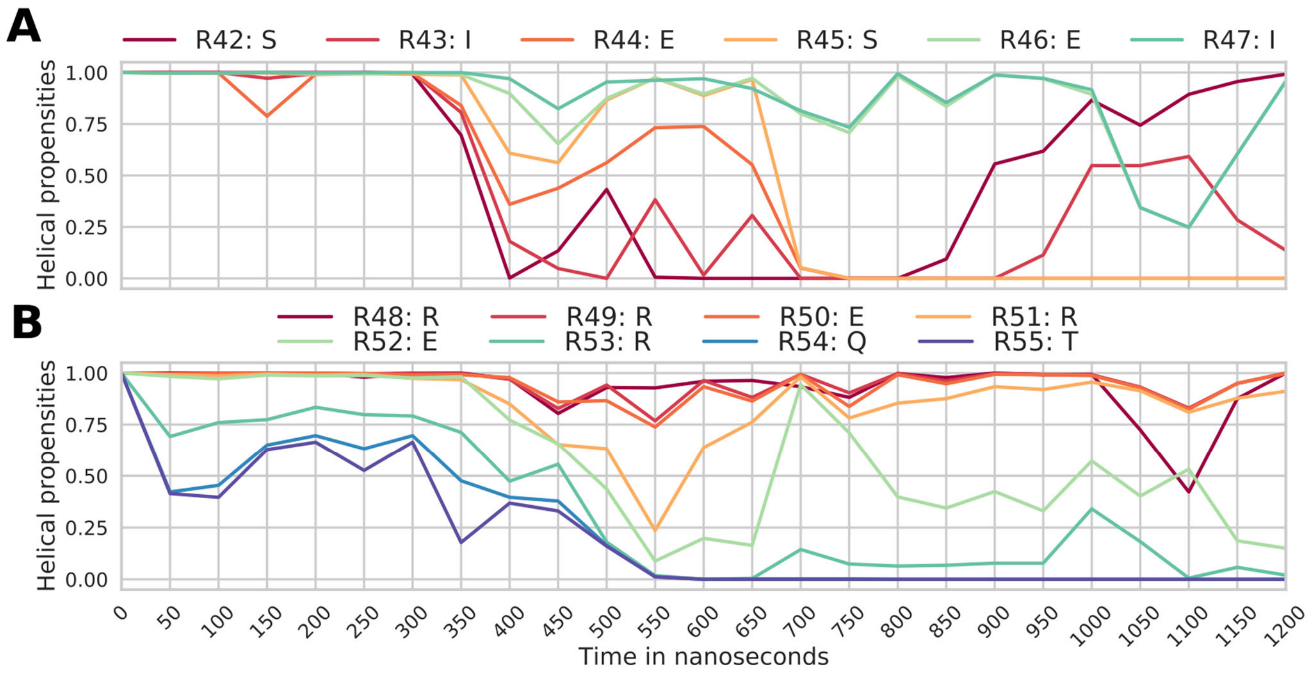


**Supplementary figure S1. Helical propensities of a section of the original TERA sequence along the simulation trajectories.**
**(A)** Helical propensity per residue for the six residues previous to TERA polyER (SIESEI), and **(B)** the six residues of the polyER and 2 residues following (RRERERQT).

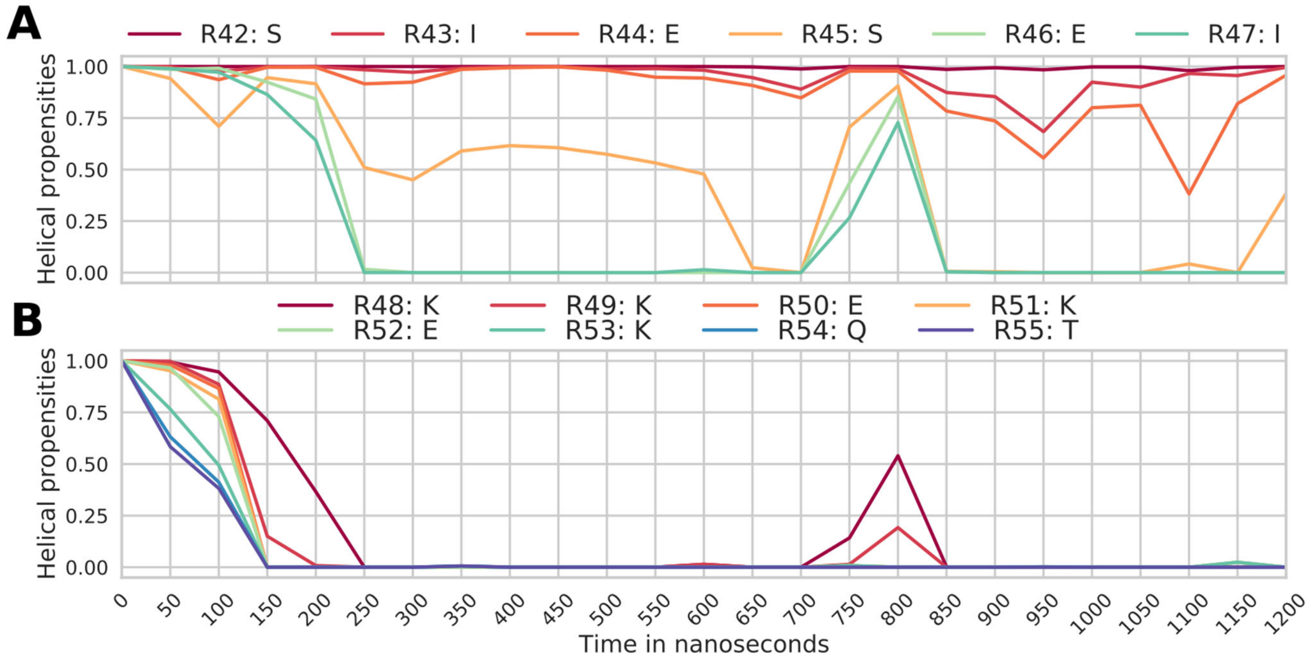

**Supplementary figure S2. Helical propensities of a section of the mutated KKEKEK along the simulation trajectories.**

**(A)** Helical propensity per residue for the six residues previous to the mutated polyEK region (SIESEI), and **(B)** the six residues of the mutated polyEK region and 2 residues following (KKEKEKQT).

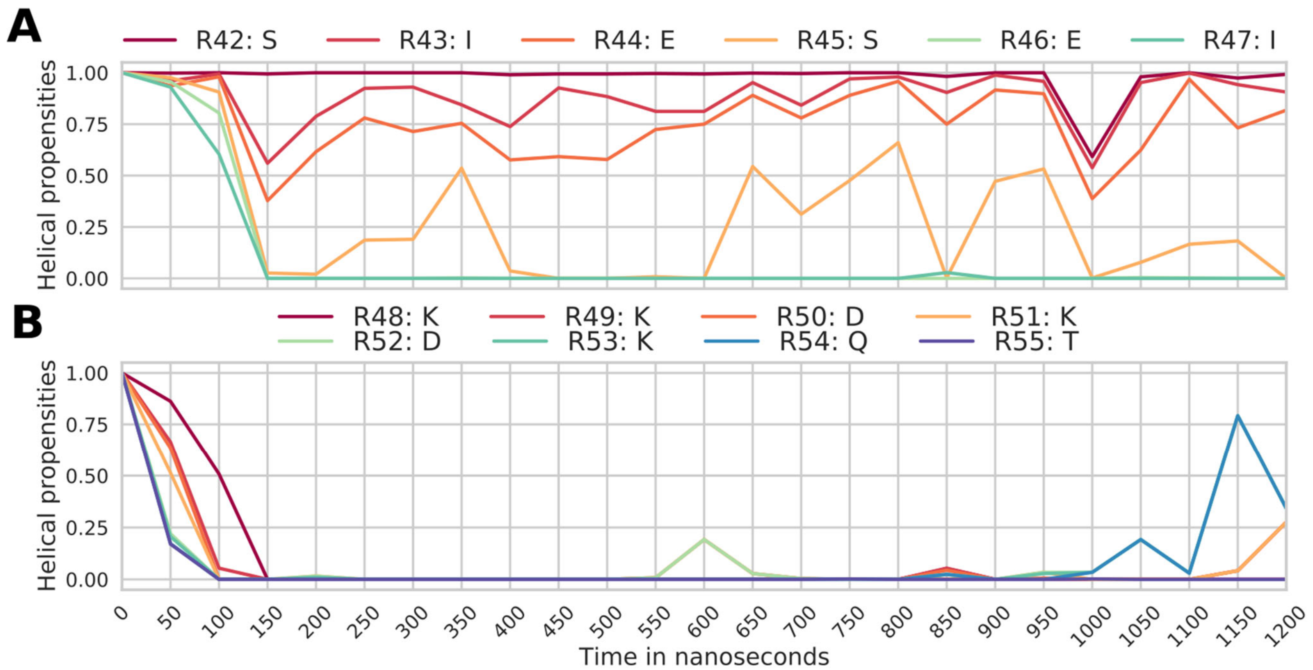


**Supplementary figure S3. Helical propensities of a section of the mutated KKDKDK along the simulation trajectories.**

**(A)** Helical propensity per residue for the six residues previous to the mutated polyDK region (SIESEI), and **(B)** the six residues of the mutated polyDK region and 2 residues following (KKDKDKQT).

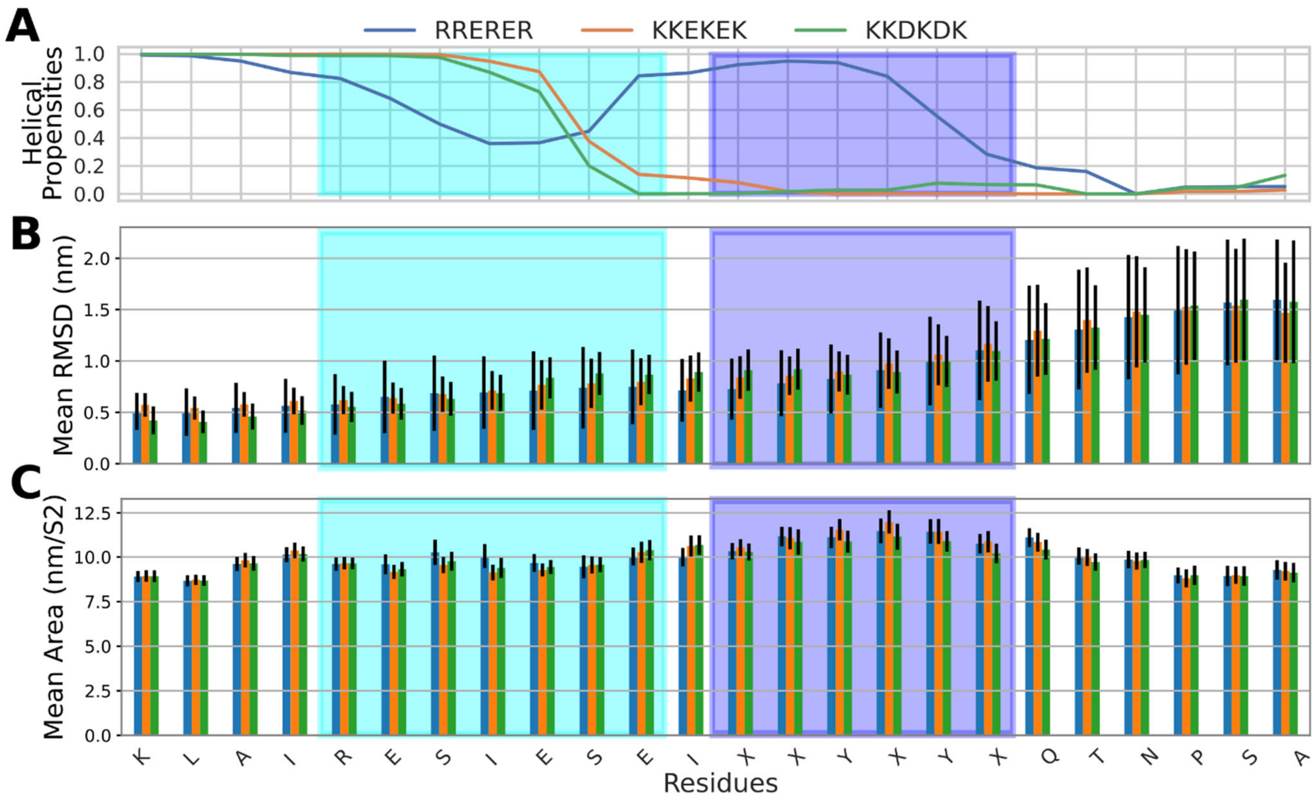


**Supplementary figure S4. RMSD and SASA of close neighbors of the polyXY region.**

**(A)** Helical propensity per residue in positions 36 to 59 of the 100-residue TERA fragment subjected to MD simulation. **(B)** Average RMSD for a six amino acid region surrounding each residue, with standard deviations. **(C)** Average area of SASA, with standard deviations. Data is shown for original polyER, RREREE (blue), and for mutated sequences: KKEKEK (orange) and KKDKK (green). The cyan box delimits the region with the largest decrease in helical propensities for the original polyER sequence, and the purple box delimits the polyXY region.

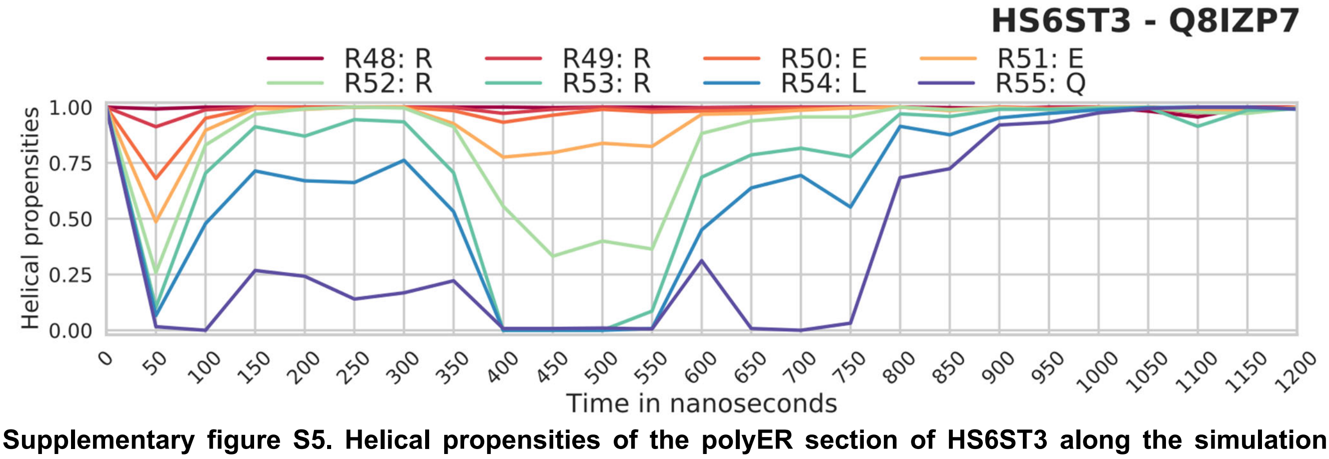


**Supplementary figure S5. Helical propensities of the polyER section of HS6ST3 along the simulation trajectories.**

Helical propensity per residue for the 6 residues of the polyER region of HS6ST3 and two following residues (RREERRLQ).

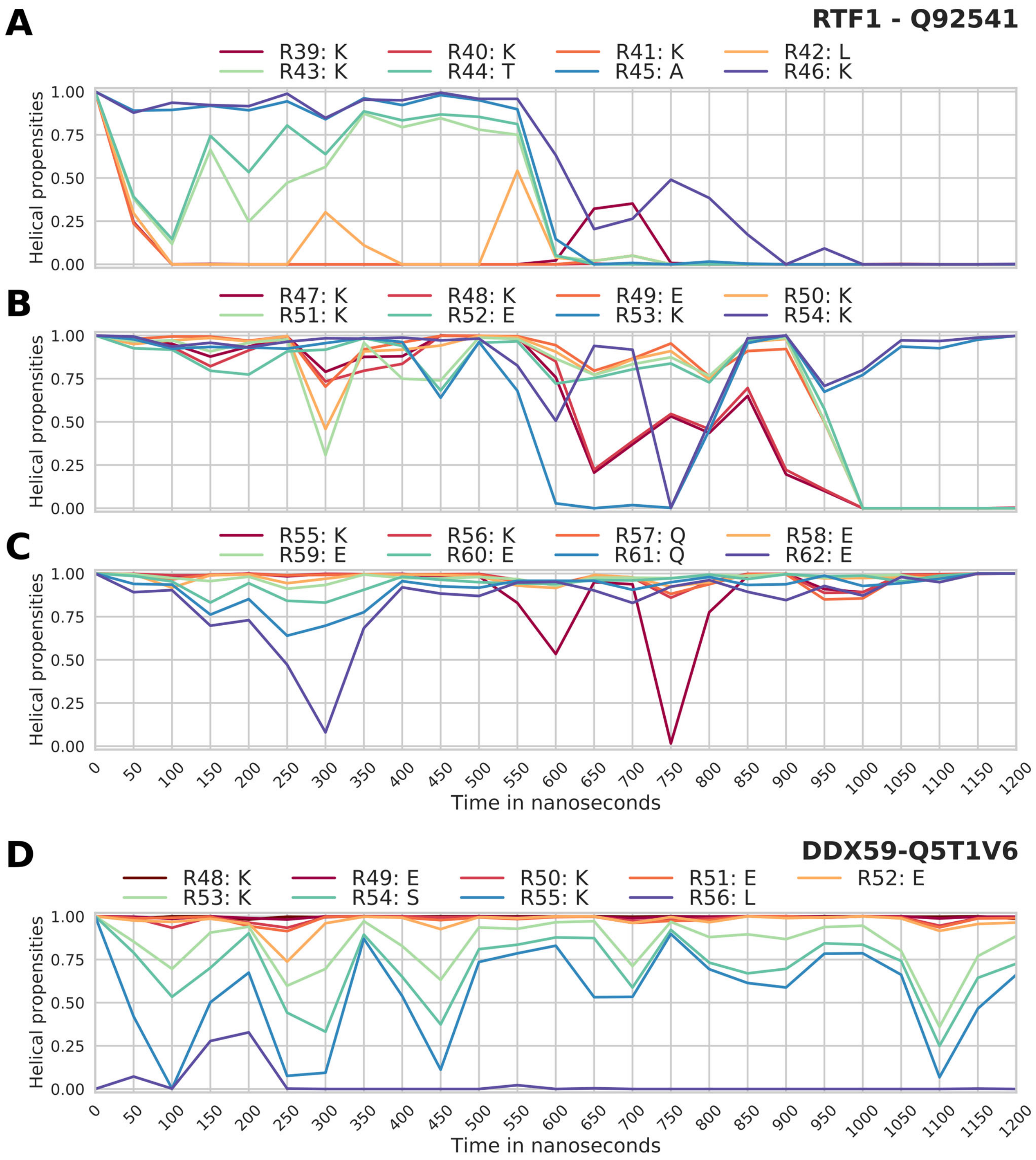


**Supplementary figure S6. Helical propensities of sections of RTF1 and DDX59 along the simulation trajectories.**

Helical propensity per residue of RTF1 **(A)** for eight residues before the polyEK region (KKKLKTAK); **(B)** the polyEK region itself (KKEKKEKK); and **(C)** the following eight residues (KKQEEEQE) of RTF1. **(D)** Helical propensity per residue of DDX59 for the six residues of the polyEK region and adjacent three residues (KEKEEKSKL).

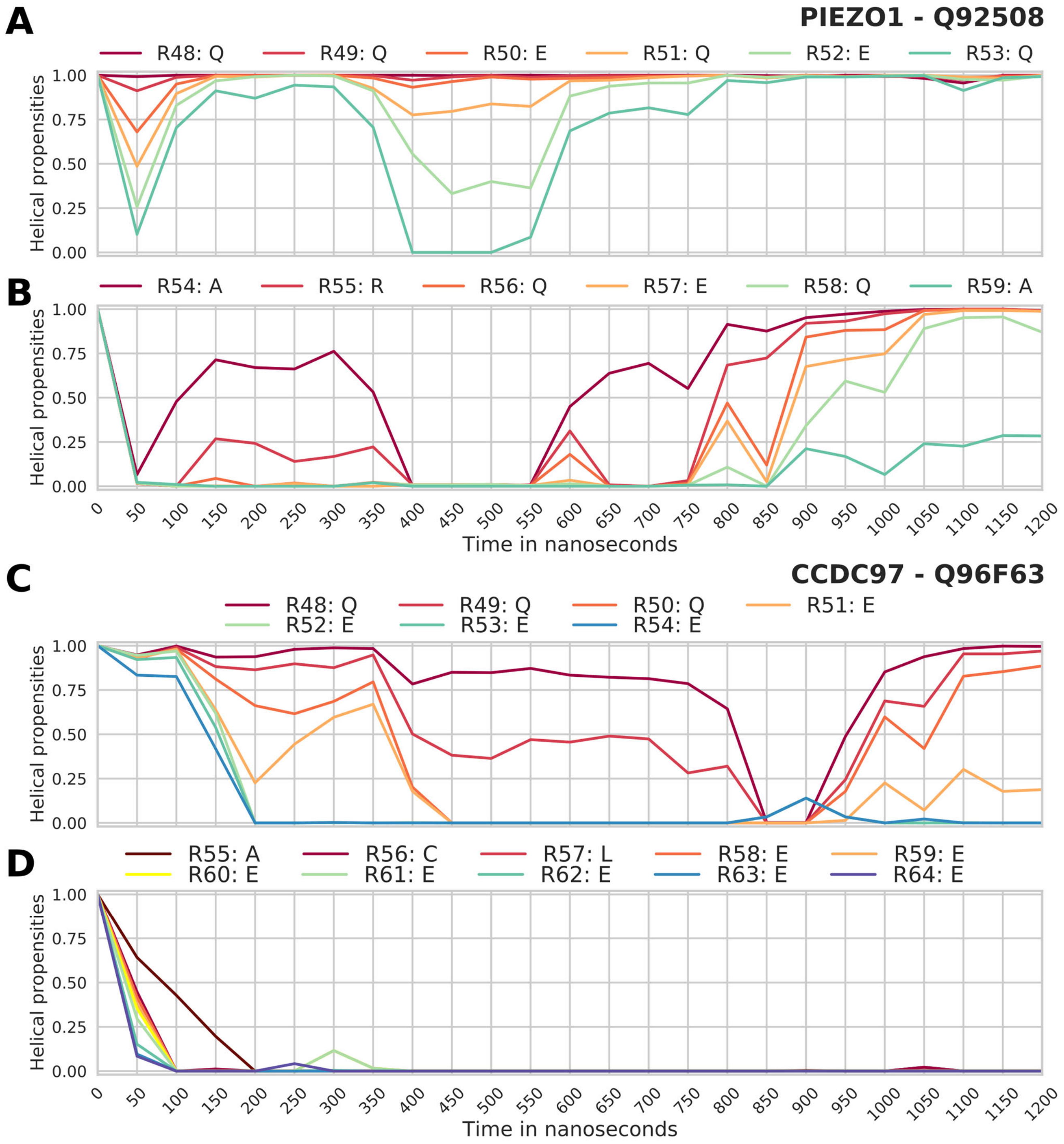


**Supplementary figure S7. Helical propensities of sections of PIEZO1 and CCDC97 along the simulation trajectories.**

Helical propensity per residue for PIEZO1 **(A)** of the polyEK region itself (QQEQEQ); and **(B)** the following six residues (ARQEQA) of PIEZO1. Helical propensity per residue for CCDC97 **(C)** of the seven residues of the polyEQ; and **(D)** ten adjacent residues (ACLEEEEEEE).

### Supplementary Tables

**Supplementary table S1. Detailed list of all human IDRs containing polyXYs and their properties.**
Columns indicate: UniProt identifier (seq name), identifier of the polyXY (poly name), amino acid XY, start and end of the polyXY in the query sequence (poly start and poly end), polyXY size, start and end of the IDR in the query sequence (idr start and idr end), size of the IDR (idr size), sequence of the polyXY (poly aa), simplified secondary structure from DSSP in the polyXY including three structure types (H: helix, E: sheet and O: Others) and blanks (poly ss), polyXY median plddt, sequence of the IDR (idr aa), simplified secondary structure from DSSP in the IDR including three structure types (H: helix, E: sheet and O: Others) and blanks (idr ss), 50 amino acid sequence to the left and to the right of the polyXY center (poly seq left and poly seq right) and their simplified secondary structure annotations (poly ss simplified left, poly ss simplified right), with pipe signs ("|||") indicating the end of the sequence range, information of the presence of a N-terminal helix (idr N-terminal), and of a C-terminal helix (idr C-terminal).

**Supplementary table S2. Subset of polyXYs within IDRs with a helical N-terminal.**
Columns indicate: Identifier of the polyXY (poly name), amino acid XY, indication of polyXY group regarding helical coverage with two different groups: "high helical overlap" and "low/no helical overlap" (poly group), median pLDDT ranged by group (median plddt), grouped distance between polyXY center and the end of the closest N-terminal helix (distance helix), grouped distance between the polyXY center and the closest previous domain (distance domain), coded pLDDT values for the 50 amino acid sequence to the left and to the right of the polyXY center (according to AlphaFold 2 specifications: D: Very low (pLDDT < 50); C: Low (70 > pLDDT > 50); B: High (90 > pLDDT > 70); and A: Very high (pLDDT > 90); coded plddt left and coded plddt right), whether the polyXY is covered by the same helix extended from the IDR end (same helix), coordinates of the targeted helix in the query sequence (helices coords), closest previous PFAM domain, when available (pfam), and its coordinates in the query sequence (pfam coords).

**Supplementary table S3. Subset of polyXYs within IDRs with a helical C-terminal.**
Columns indicate: Identifier of the polyXY (poly name), amino acid XY, indication of polyXY group regarding helical coverage with two different groups: "high helical overlap" and "low/no helical overlap" (poly group), median pLDDT ranged by group (median plddt), grouped distance between polyXY center and the start of the closest C-terminal helix (distance helix), grouped distance between the polyXY center and the closest following domain (distance domain), 50 letter coded pLDDT values for the 50 amino acid sequence to the left and to the right of the polyXY center (according to AlphaFold 2 specifications: D: Very low (pLDDT < 50); C: Low (70 > pLDDT > 50); B: High (90 > pLDDT > 70); and A: Very high (pLDDT > 90); coded plddt left and coded plddt right), whether the polyXY is covered by the same helix extended from the IDR end (same helix), coordinates of the targeted helix in the query sequence (helices coords), closest following PFAM domain, when available (pfam), and its coordinates in the query sequence (pfam coords).

**Supplementary table S4. Sequence details of fragments submitted to MD simulations.**

Sequence fragment, initial secondary structure annotations and median pLDDT of the polyXY region of AlphaFold2 predicted regions submitted to MD simulations.

| Gene / UniProt | * Coded pLDDT<br>Original sequence<br>Sequence composition<br>Secondary structure composition at 1.2 µs | ** Median pLDDT |
|---|---|---|
| TERA<br>P55072 | AABAAAAAAAAAAAAAAAAAAAAAAAAAAAAAAAAAAAAAAAABABBBBBBBBBBBBCCCCCCCCCBBBBAAAAAAAAAAAABBBABBABBBBBBBBBBBB<br>LRKSPVAKDVDLEFLAKMTNGFSGADLTEICQRACKLAIRESIESEIRRERERQTNPSAMEVEEDDPVPEIRRDHFEEAMRFARRSVSDNDIRKYEMFAQ<br>HOOO E OO HHHHHHHOOO HHHHHHHHHHHHHHHHHHHHHHHHHHHHHHHH OOO O OE HHHHHHHHHO OO HHHHHHHHHHHH | 81.13 |
| HS6ST3<br>Q8IZP7 | BBAAAAAAAAAAAAAAAAAAAAAAAAAAAAAAAAAAAAAAAAAAAAAAAAAAABABBBBCCDDDDDDDDDDDDCCCDCCDCCDD<br>VEINEGARQRIEDLNFLDMQLYEYAKDLFQQRYHHTKQLEHQRDRQKRREERRLQREHRDHQWPKEDGAAEGTVTEDYNSQVVRW<br> HHHHHHHHHHOHHHHHHHHHHHHHHHHHHHHHHHHHHHHHHHHHHHHHHHHHHHHO HHHHHHH | 93.36 |
| RTF1<br>Q92541 | BBBBBBBBBBBBBBBBBBBBBBBBBBBBABBBBBBBBBBBBBBBBBBBBBBBBBBBBBCCCCCCCCCCDDDDDDDDCCCCCCCCCCCCCBBBBBBBBBB<br>MGDEEDRARLEQMTEKEREQELFNRIEKREVLKRRFEIKKKLKTAKKKEKKEKKKKQEEEQEKKKLTQIQESQVTSHNKERRSKRDEKLDKKSQAMEELK<br> OHHHHHHHHHO HHHHHHHHHHHHHHHHHHHHHHHHHHHHHHHHHHHHHHHHHHHHHHHHHHHHOOOOO OOOHHHHHHHHHHHHHHHHHHHHH | 82.78 |
| DDX59<br>Q5T1V6 | BBBBBBABBBBBBBBABBABAABBBBBBBBBBBBBABABBBBBBBBBBBBBBCCCCDDDDDDDDDDDDDDDCCBBBBBBBBBBBBBBBCBBCBBBCBBBA<br>SFSKTQRWAEPGEPICVVCGRYGEYICDKTDEDVCSLECKAKHLLQVKEKEEKSKLSNPQKADSEPESPLNASYVYKEHPFILNLQEDQIENLKQQLGIL<br>OOHHHO E OOO E OOOOEE EEEOOOOEEEOOHHHHHHHHHHHHHHHHHHHH HHHO HHHHOO HHHHHHHHHHOOEE | 75.97 |
| PIEZO1<br>Q92508 | BBBBBBCCCCCCDDDDDDDDDDDDCCCCCCCCCCCCCCCCCBCBBBCBBBCBCCCCCCDDDDDDDDDDDDDDDDDDDDDDDDDDDCCCCCCBCCBBBBB<br>GDYFLFESDSEEEEEAVPEDPRPSAQSAFQLAYQAWVTNAQAVLRRRQQEQEQARQEQAGQLPTGGGPSQEVEPAEGPEEAAAGRSHVVQRVLSTAQFLW<br> HHHHOOOOO O HHHHHHHHHHO HHHHHHHHHHHHHHHHHHHHOO O HHHHHHHHHHHHHH | 71.35 |
| CCDC97<br>Q96F63 | AAABBBBBBBCCCDDDDCDCCDDCCCCBBBBBAABAAAAAAAAAAAAAAAAAAAAAABBBBBBBCCCCCDDDDDDDDDCCBBBAAAAAAAAAAAAAAAAAA<br>YLTQEELSARTPTHQPPKPGSPGRPACPLSNLLLQSYEERELQQRLLQQQEEEEACLEEEEEEEDSDEEDQRSGKDSEAWVPDSEERLILREEFTSRMHQ<br>H HHHHHHHOOO OO OO HHHHHHHHHHHHHHHHHHHHHHHHHHHHHHHHHHHOOO HHHHHHHHHHHHHHHHH | 96.30 |

* pLDDT scores letter coded according to AlphaFold 2 specifications: D: Very low (pLDDT < 50); C: Low (70 > pLDDT > 50); B: High (90 > pLDDT > 70); and A: Very high (pLDDT > 90). The bold region indicates the target helix, underlined section the neighbor domain, blue region delimits the polyXY region and red delimits the IDR region. Domains are not present in all segments.

** Median pLDDT of the polyXY region
